# Does capacity to elevate androgens underlie variation in female ornamentation and territoriality in White-shouldered Fairywren (*Malurus alboscapulatus*)?

**DOI:** 10.1101/2023.02.16.528857

**Authors:** Jordan Boersma, Erik D. Enbody, Serena Ketaloya, Heather E. Watts, Jordan Karubian, Hubert Schwabl

## Abstract

Historic bias toward study of male sex hormones and sexual signals currently constrains our perspective of hormone—behavior—phenotype relationships. Resolving how ornamented female phenotypes evolve is particularly important for understanding the diversity of social signals across taxa. Studies of both males and females in taxa with variable female phenotypes are needed to establish whether sexes share mechanisms underlying expression of signaling phenotypes and behavior. White-shouldered Fairywren (*Malurus alboscapulatus*) subspecies vary in female ornamentation, baseline circulating androgens, and response to territorial intrusion. The *moretoni* ornamented female subspecies show higher female, but lower male androgens, and a stronger pair territorial response relative to pairs from the *lorentzi* unornamented female subspecies. Here we address whether subspecific differences in female ornamentation, baseline androgens, and pair territoriality are associated with ability to elevate androgens following gonadotropin releasing hormone (GnRH) challenge and in response to simulated territorial intrusion. We find that subspecies do not differ in their capacity to circulate androgens in either sex following GnRH or territorial intrusion challenges. Whereas pre-GnRH androgens were somewhat predictive of degree of response to territorial intrusions, higher androgens were associated with lower territorial aggression. Post-GnRH androgens were not predictive of response to simulated intruders, nor did females sampled during intrusion elevate androgens relative to flushed controls, suggesting that increased androgens are not necessary for the expression of territorial defense behaviors. Collectively, our results suggest that capacity to produce and circulate androgens does not underlie subspecific patterns of female ornamentation, territoriality, and baseline androgens.

## 1. Introduction

Androgens mediate a variety of morphological and behavioral traits across diverse vertebrate taxa (Cox et al., 2016; Hau, 2007; Hau and Goymann, 2015). For instance, in the context of breeding, elevated androgens in male birds can cause both molt into ornamental nuptial plumage (Kimball and Ligon, 1999; Lindsay et al., 2009; Peters et al., 2000) and expression of territorial and sexual behavior at the onset of breeding (Day et al., 2006; Gleason et al., 2009; Goymann, 2009; Raouf et al., 1997; Schwabl et al., 2005; Wiley and Goldizen, 2003; Wingfield et al., 1990). These pleiotropic actions of androgens can link morphology and behavior to produce variation in integrated phenotypes (Lipshutz et al., 2019). Comparative androgen studies among individuals or populations varying phenotypically can help identify which phenotypic traits might be under common androgenic control. Due to the transient nature of hormone production and release and the myriad causes of rapid elevations in circulating levels (Goymann, 2009; Oliveira, 2004), it can, however, be difficult to draw conclusions about phenotypic integration from single hormone samples (Cox et al., 2016). Gonadotropin-releasing hormone (GnRH) causes a trophic cascade that stimulates the gonads to produce and release androgens (Bergeon Burns et al., 2014a; McGlothlin et al., 2008a). Intramuscular injections of GnRH (“GnRH challenges”) produce repeatable measurements of an individual’s capacity to produce androgens, thus providing a reliable assessment of potential constraints on androgen-regulated signaling (Ambardar and Grindstaff, 2017; Bergeon Burns et al., 2014a; Cain and Pryke, 2017; Jawor et al., 2007; McGlothlin et al., 2007).

Differences in secretion of androgens in response to GnRH challenge have been shown to correlate with integrated morphological and behavioral phenotypes in some male birds, suggesting that varying capacity to circulate androgens underlies phenotype expression (Cain and Pryke, 2017; McGlothlin et al., 2008b; Mills et al., 2008; Spinney et al., 2006). GnRH challenge studies in females that vary phenotypically are limited, with female White-throated Sparrow (*Zonotrichia albicolis*) morphs showing no difference in GnRH-induced testosterone levels, despite morph-specific variation in male GnRH response in the same species (Spinney et al., 2006), and a study of Dark-eyed Junco (*Junco hyemalis*) finding that response to GnRH might underlie differences in reproductive timing across populations (Kimmitt et al., 2020). Despite the commonality of female ornamentation in nature, there has been relatively little research on its mechanistic basis, leaving a gap in our understanding of the processes of signal evolution. Androgens appear to mediate female ornamentation in some polyandrous bird species (Johns, 1964; Muck and Goymann, 2011), though studies in more common mating systems are sparse (Lahaye et al., 2014; Moreno et al., 2014). The avian ovary synthesizes and secretes androgens and can be stimulated by GnRH injection to do so (Bentz et al., 2022; Egbert et al., 2013; Ketterson et al., 2005; Tilly et al., 1991), although circulating levels of androgens, including testosterone, are generally lower in females than males (Goymann and Wingfield, 2014; Ketterson et al., 2005; Møller, A. P., Garamszegi, L. Z., Gil, D., Hurtrez-Boussè, S., Eens, 2005). Furthermore, exogenous testosterone can stimulate molt of ornamented feathers in otherwise cryptically-colored females (Boersma et al., 2020; Lank et al., 1999; Lindsay et al., 2016). Thus, female birds of at least some species have the capacity to produce androgens and have the necessary physiological response mechanisms (i.e. receptors and genes) to develop ornamentation, even when it is not developed naturally. Variable capacity to produce androgens in response to GnRH is therefore a prime candidate for explaining diversity in female ornamentation.

Until recently, signaling traits like ornaments were understudied in females and were thought to be simply the byproduct of selection on male elaboration (Darwin, 1871; Lande, 1980). Yet female ornaments appear to often function in the context of competition for ecological resources (i.e. social selection, Tobias et al., 2012), though they can also serve to attract mates (i.e. sexual selection, (Hare and Simmons, 2019; Joseph A Tobias et al., 2012). Territorial defense is ubiquitous across avian taxa, and both sexes often defend territories jointly as breeding pairs (Burtka and Grindstaff, 2015; Cain et al., 2015). Androgens have been shown to correlate with and stimulate expression of territory defense behaviors in many male (Goymann et al., 2015; Hau and Goymann, 2015; Oliveira et al., 2009), and some female bird species (Rosvall, 2013).

That androgens often increase in males during periods of enhanced territory competition led to the formulation of the challenge hypothesis, which posits that conflict with same sex rivals causes rapid elevations of androgens (Wingfield et al., 1990).

Evidence for agonistic encounter induced androgen elevations is mixed (reviewed in (Goymann et al., 2019; Hirschenhauser and Oliveira, 2006), though most studies are limited to males, despite the ubiquity of joint territorial defense (Hau et al., 2000; Horton et al., 2012; Rosvall, 2013; Schwabl et al., 2005).

In order to tease apart whether low androgens measured during an agonistic encounter are due to low due to limited physiological capacity or because the stimulus does not elicit maximal androgen release, one can sample androgens during territory defense and again after a GnRH challenge (Apfelbeck and Goymann, 2011). In other words, androgen levels that appear low compared to other individuals can be maximal for that individual as determined by GnRH challenge due to that individual having a lower capacity to produce androgens. Establishing what maximal androgen levels are for each individual allows one to determine whether a given stimulus causes elevation of androgens beyond baseline.

Here we assess whether capacity to produce androgens underlies phenotypic differences in female ornamentation and territoriality across subspecies of White-shouldered Fairywren (*Malurus alboscapulatus*). The *moretoni* subspecies of this New Guinea endemic bird species is characterized by ornamented females and greater pair territoriality, whereas the females of the *lorentzi* subspecies are cryptic brown (‘unornamented’) and pairs defend territories less aggressively and show greater social connectivity (Boersma et al., 2022; Enbody et al., 2018). In addition to differences in female ornamentation and social behavior, the subspecies also differ in their baseline circulating androgen levels. Plasma androgens are higher on average in ornamented *moretoni* females relative to unornamented *lorentzi* females (Enbody et al., 2018); however, male androgens are higher on average in *lorentzi* compared to *moretoni* despite males being ornamented in both subspecies (Boersma et al. 2022). A testosterone-implant experiment with unornamented *lorentzi* females revealed that androgens seem to be linked to development of some female ornamentation (white shoulder patch), which then enhanced territory defense (Boersma et al., 2020). Finally, an experimental study found that male androgens increased subtly in response to simulated territorial intrusions relative to flushed controls, however levels were much higher when sampled during courtship competition (Boersma et al., 2022a).

Collectively, previous studies of White-shouldered Fairywren have shown that androgens are linked to the development of ornamentation in females and territorial and sexual behavior in males (Table 1). In the current study, we use GnRH and simulated territorial intrusion challenges to assess whether varying capacity to produce and circulate androgens underlies subspecific variation in female ornamentation, territorial behavior, and baseline androgens. In conjunction with a recent study of male androgens (Boersma et al., 2022a), we also test whether simulated territorial intrusions cause females to elevate androgens relative to flushed controls (“challenge hypothesis” in females; Wingfield et al.,1990).

**Table 1.**
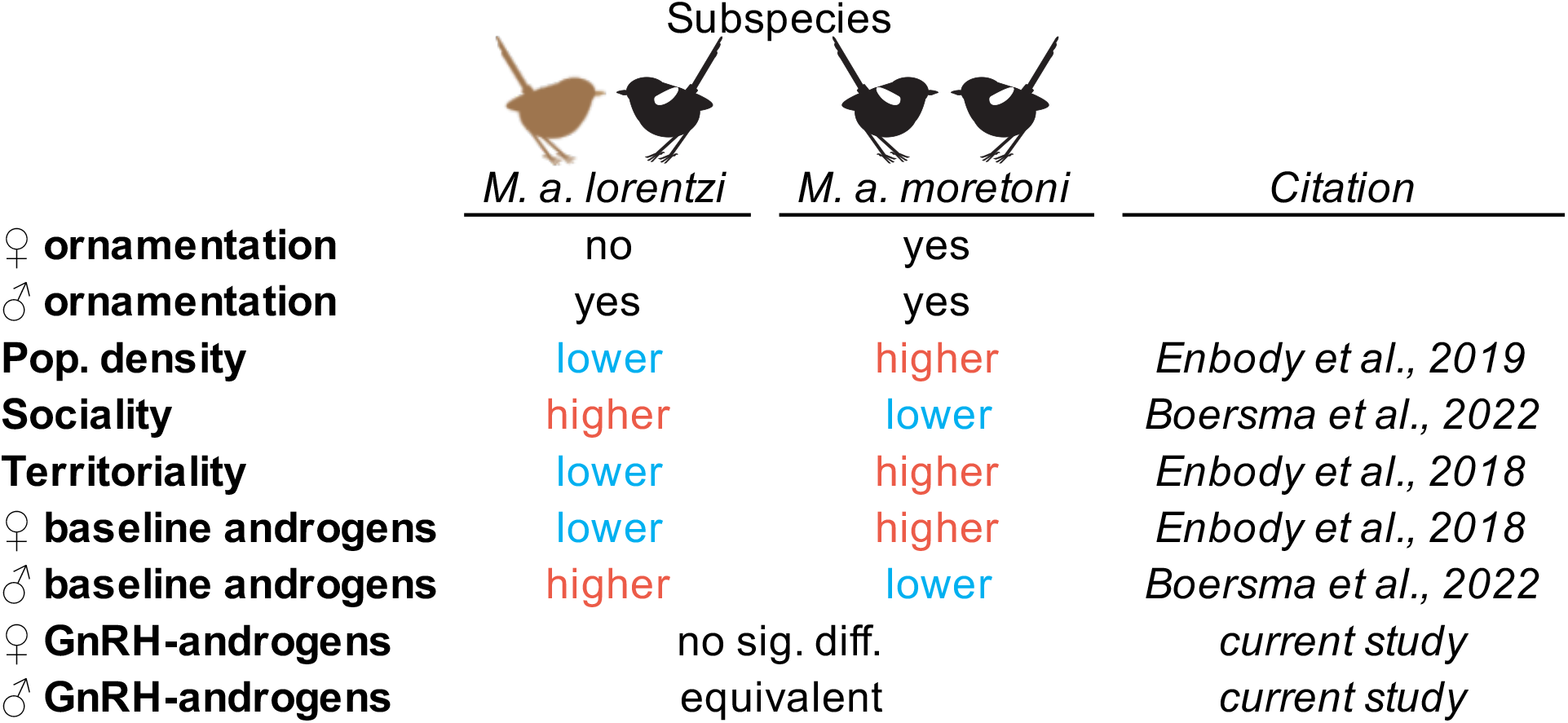
Comparison of subspecies in plumage ornamentation, social environment and behavior, and androgen circulation. Citations are listed for each comparative study.

|  | Subspecies |  |  |
| --- | --- | --- | --- |
|  | *M. a. lorentzi* | *M. a. moretoni* |  |
| ♀ ornamentation | no | yes |  |
| ♂ ornamentation | yes | yes |  |
| Pop. density | lower | higher | *Enbody et al., 2019* |
| Sociality | higher | lower | *Boersma et al., 2022* |
| Territoriality | lower | higher | *Enbody et al., 2018* |
| ♀ baseline androgens | lower | higher | *Enbody et al., 2018* |
| ♂ baseline androgens | higher | lower | *Boersma et al., 2022* |
| ♀ GnRH-androgens | no sig. diff. |  | *current study* |
| ♂ GnRH-androgens | equivalent |  | *current study* |

## 2. Materialized and Methods

### 2.1 Study system and general field methods

White-shouldered Fairywrens are endemic to savanna and edge habitat in New Guinea and belong to the avian family Maluridae (Schodde, 1982). This species shows low levels of cooperative breeding, and pairs defend territories and can breed year-round (Enbody et al., 2019, 2018). Females alone build nests and incubate while males provision females during these stages and then assist females in feeding nestlings (Enbody et al., 2019). We studied populations belonging to the *moretoni* subspecies which contains females with contrasting black-and-white (‘ornamented’) plumage akin to males, and populations of the *lorentzi* subspecies, containing cryptic brown (‘unornamented’) females in Papua New Guinea. Our *moretoni* work was conducted around the villages of Podagha (149º 90′ E, 9º69’ S, 50-60 m) and Porotona, Milne Bay Province (150° 35′ E, 10°15′ S, 10–20 m a.s.l.). We conducted *lorentzi* work in two populations near Obo village, Western Province: one was along the western edge of the Fly River drainage (141°19′ E, 7°35′ S, 10–20 m a.s.l.), and the other was a chain of islands in the Fly River (141°16′ E, 7°34′ S, 10–20 m a.s.l.).

We captured individuals in mistnets between 6:15 – 11:10 and 16:16 – 18:00 during peaks of daily Fairywren activity from May 23 to July 24, 2019. Pairs were randomly assigned to be flushed into nets absent a social stimulus or during a simulated territorial intrusion (see full details in section 2.2). In conjunction with a previous experiment (Boersma et al., 2022a), some *lorentzi* males (N = 14) were captured during extra-pair courtship competitions. We assessed breeding stage based on whether pairs had recently fledged young with them, and whether captured females had a distended abdomen indicative of egg laying or had a defeathered brood patch. For females with brood patches, but no distended abdomen, we used degree of vascularization and wrinkling of skin to estimate whether pairs were in the incubation or nestling stage. Pairs containing a female carrying an egg and/or exhibiting a highly vascularized brood patch were grouped in stage 1 (laying and incubating); pairs with fledglings and/or a female with a wrinkly brood patch were grouped in stage 2 (nestling and fledgling); and stage 3 contained all pairs with no signs of breeding activity and cases of males captured absent their mate where we were unable to estimate stage (N = 13 males of unknown stage). Capture, blood sampling, GnRH injection, and behavioral protocols were all approved by the Institutional Animal Care and Use Committee (IACUC) at Washington State University (protocol #: ASAF-04573).

### 2.2 Experimental design

#### 2.2.1 GnRH challenges

We collected blood samples from the jugular vein for each individual at two time-points, once immediately (i.e., within 10 minutes) after capture in a mistnet to assess pre-GnRH androgens and again after ~30 minutes (range: 29.6 – 41.2 min, mean = 32.0 min) following administration of gonadotropin-releasing hormone (GnRH). GnRH challenges consisted of a standard dose of 500 ng of chicken GnRH-I (Bachem H-3106, Torrance, CA, USA) dissolved in 10 μl of sterile saline injected into the left pectoralis major. This dose was determined based on previous GnRH work in a congener (Barron et al., 2015) and another similarly-sized passerine (McGlothlin et al., 2008b). All birds were held in a cloth bag between initial and post-GnRH sample collection. We aimed for ~20 μl of plasma at both time points; pre-GnRH samples ranged from 12.1 – 32.6 μl (mean: 23.0 μl) and post-GnRH samples from 8.4 – 33.1 μl (mean: 23.2 μl).

#### 2.2.2 Androgens during simulated territorial intrusions

In conjunction with a companion study (Boersma et al. 2022), pairs were captured while actively defending territories against simulated intruders to test whether pre-GnRH or post-GnRH androgen titers explained intensity of territorial defense. We used an established simulated territorial intrusion protocol (STI) meant to mimic territorial disputes in this species (full details in Boersma et al. 2022, Enbody et al. 2018). Briefly, cardstock mount pairs of the focal subspecies (N = 4 exemplars for each sex) were placed within the territory of a focal pair and recorded duet song (N = 8 exemplars of local subspecies) was played from a nearby speaker to elicit territory defense. To assess territorial defense, we recorded latency and rate of solo songs/pair duets (combined) and flybys, which occur when an individual flies within 2m of mounts, as well as latency and proportion of time within 5m of mounts. We quantified these behaviors for 10 minutes, at which point we opened a furled net that was placed approx. 5m from mounts. We did not observe any effect of the furled net on the behavior of territory defending pairs, and all individuals were captured while responding to the intrusion. The duration of playback until capture ranged from 11 – 68 min (mean = 17.84 min) for females and 11 – 43 min (mean = 17.71 min) for males. In all cases individuals included in this experiment had an initial blood sample taken within 10 minutes of capture (mean = 4.09 min for females and 4.17 min for males). Following the initial blood sample, we injected individuals with GnRH (same details as above) and collected another blood sample ~30 min. later (range: 29.47 – 41.47 min).

#### 2.2.3 Challenge hypothesis experiment in females

To test whether females elevate androgens during territorial defense, we compared females caught while defending territories (using the simulated territorial intrusion protocol described above) versus flushed controls. Flushed controls were females that were foraging on their territory absent conspecifics outside of their familial group at the time of capture. We used slow movements to guide individual pairs into mistnets to avoid eliciting a stress response. All blood samples were taken within 8 minutes of capture (mean = 4.15 min; range: 2.08 – 7.5 min).

### 2.3 Plasma storage and radioimmunoassay

At the end of each field day, we centrifuged blood samples to separate plasma from red blood cells and used a Hamilton™ syringe to transfer plasma into 0.4 ml of 200 proof ethanol (Fisher Bioreagants™) in individual Eppendorf™ tubes. Samples were stored at room temperature in the field, then transferred to a refrigerator in our lab at Washington State University. Prior to assay we randomized samples across assays to account for inter-assay variation. We used an established radioimmunoassay protocol to measure total androgen titres (Barron et al., 2013; Lantz et al., 2017; Lindsay et al., 2011). Total androgens are reported instead of testosterone specifically because the antibody we used (Wien Laboratories T-3003, Flanders, NJ, USA) cross-reacts strongly with 5α-dihydrotestosterone (DHT). Our minimum detectable androgen concentration of 445.45 pg/ ml based on 8.4 µl (lowest in this study) plasma volume with a mean recovery rate of 65.77%. We used our assay’s minimal detectable level of 1.95 pg/ tube to calculate and assign a total androgen concentration to undetectable samples (N = 53 out of 191 total samples).The inter-assay coefficient of variation across 5 assays was 13.57% (calculated following methods in (Chard, 1995).

### 2.4 Statistical methods

All analyses were conducted in R (<www.r-project.org>) version 3.6.1. We natural log transformed both pre-GnRH and GnRH-induced androgens prior to analysis. We compared subspecies in response to GnRH using linear models, binomial generalized linear models, and ANOVA in base R (R Core Team, 2019). We used backward stepwise selection throughout to remove candidate predictors with *p* ≥ 0.2 (following Wang et al., 2008). We confirmed heteroscedasticity and inspected Q-Q residual plots to check for approximate normality for our linear models. When group comparisons were null (i.e. GnRH-induced androgen across subspecies and baseline androgens across female sampling contexts) we used equivalence tests in R package TOSTER (Lakens, 2017) to test whether means were equivalent. We used this approach to test whether null comparisons across groups were indicative of no meaningful difference in androgens rather than a constraint of small sample sizes obscuring our ability to detect differences. For GnRH-induced androgens we set equivalence bounds according to mean differences across subspecies previously measured in females (Enbody et al., 2018) and males (Boersma et al., 2022a). For simulated territorial intrusion versus flushed androgen sampling comparison in females, we used the previously measured mean difference in males across these same sampling contexts as equivalence bounds (Boersma et al., 2022a).

#### 2.4.1 Response to GnRH challenges across subspecies

We ran separate models for males and females comparing subspecies in three metrics of response to GnRH: a) binary response (Y/N) variable which reflects whether androgens increased from pre-GnRH to post-GnRH sampling regardless of magnitude of increase, b) change (Δ) in log-transformed androgens from pre-GnRH to post-GnRH sampling, and c) GnRH-induced log-transformed androgen levels. For the latter, we only analyzed individuals who increased androgens following GnRH challenge as we were interested in comparing subspecies without the confound of individuals who were physiologically incapable of responding to GnRH. Breeding stage, time of day bled, and delay time between GnRH injection and second blood sample were included as covariates for all three response variables. We removed three outliers (two *lorentzi* males, one *moretoni* female) from Δandrogens and post-GnRH androgen analysis due to levels exceeding suspected physiological limits in this system (i.e. likely the product of measurement error). For individuals captured multiple times for this experiment we took the first sample collected (N = 1 female and 5 males).

#### 2.4.2 Androgen predictors of response to STI

We reduced our 5 simulated territorial intrusion response variables to 2 using principal component analysis (PCA) with the prcomp command. We then analyzed androgen predictors of each principal component using linear models. Males and females were run in separate PCAs to allow for sex-specific differences in response, and we analyzed principal components with eigenvalues > 1.0 in separate linear mixed models by sex. We first tested whether playback time, time-of-day bled, net-to-bleed time, and GnRH- to-bleed time affected androgen levels in this dataset; only terms with p < 0.2 were included in models. We used pre-GnRH and post-GnRH androgen variables as described above as candidate predictors with breeding stage, subspecies, and the interaction between subspecies and both androgen variables.

#### 2.4.3 Challenge hypothesis test in females

We analyzed the binary (Y/N) variable reflecting pre-GnRH androgen detectability (see section 2.3 for details) and log-transformed pre-GnRH androgen levels among females captured during STIs versus flushed controls. For both models we included breeding stage, playback time, net-to-bleed time and time-of-day bled, as well as subspecies and the interaction between subspecies and sampling context (STI vs. flush) as initial candidate predictors.

## 3. Results

### 3.1 Response to GnRH challenges across subspecies

We analyzed response to GnRH in 54 females (N = 22 *lorentzi* and 32 *moretoni*). The proportion of females who elevated androgens in response to GnRH injection did not differ among subspecies, with 5 of 22 (23%) *lorentzi* and 10 of 31 *moretoni* (32%) females elevating androgens following GnRH (Table S1a). GnRH-to-bleed time was predictive of female response to GnRH, with later times being associated with an increased probability of post-GnRH increase in androgens (*p* = 0.02; mean = 33.09 min for female’s that elevated androgens vs. 31.68 mean min for females who did not elevate androgens post-GnRH; Figure S1a). Breeding stage and time of day did not predict whether females elevated androgens following GnRH injection. Female subspecies also did not differ in Δ androgens post-GnRH (Table S1b), or GnRH-induced androgen levels (Figure 1, Table S1c). The only significant covariate for the two continuous post-GnRH variables was time-of-day bled in the GnRH-induced androgen level analysis (*p* = 0.004, Figure S1b). We failed to reject the hypothesis that mean post-GnRH androgens were equivalent across female subspecies (*t* = 0.16, *p* = 0.44; mean ± sd: 690.41 ± 525.39 pg/ml *lorentzi* vs. 1042.34 ± 801.03 pg/ml *moretoni* females).

**Figure 1.**
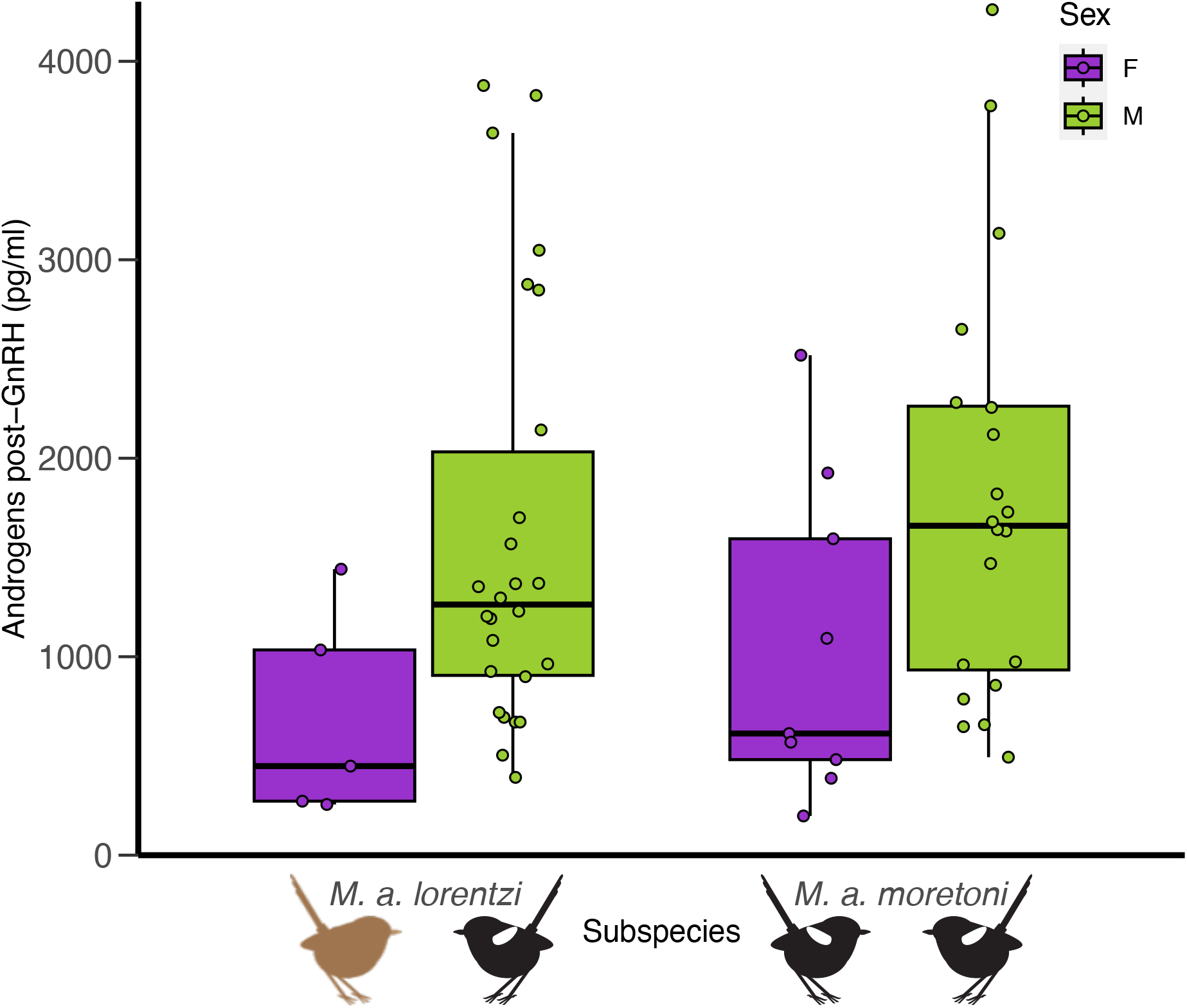
Plasma androgens following GnRH challenge among subspecies (split by sex) that differ in female ornamentation. There was no difference between subspecies among females or among males. We show only individuals who elevated androgens from initial sampling following GnRH challenge.e

Male response to GnRH was analyzed for 68 males, with 28 of 36 (78%) *lorentzi* and 20 of 32 (63%) *moretoni* elevating androgens post-GnRH. There were no statistically significant predictors of whether males increased androgens following GnRH challenge, Δ androgens post-GnRH, or GnRH-induced androgen titres (Table S2; N = 45 males who increased androgens following GnRH challenge). Mean post-GnRH androgens were equivalent across male subspecies (*t* = 2.16, *p* = 0.02; mean ± sd: 1617.75 ± 1062.28 pg/ml *lorentzi* vs. 1791.08 ± 1046.89 pg/ml *moretoni* males).

### 3.2 Androgen predictors of response to STI

We analyzed androgen predictors of the top 2 principal components (PCs) of STI response in 22 females (8 *lorentzi* and 16 *moretoni*) and 17 males (6 *lorentzi* and 11 *moretoni*). We reversed the sign of both male component scores so that high scores for both components are consistent with female PCs and generally reflect greater territory defense behavior (Table 2). High PC 1 scores in both sexes reflect rapid approaches to mounts and sustained close proximity with frequent close flights past mounts, but less vocal territoriality (i.e. longer latency and lower rate of singing/duetting). Whereas high PC 2 scores are indicative of shorter latency and more frequent songs as well as shorter latency and longer time within 5m of mounts.

**Table 2.** Principal Component Analysis (PCA) parameters and variable loadings for response to simulated territorial intrusion. We analyzed the top two principal components for **(a)** females and **(b)** males separately.

| **(a) Female Principal Component Analysis** |  |  |
| --- | --- | --- |
|  | PC1 | PC2 |
| Standard deviation | 1.73 | 1.56 |
| Proportion of Variance | 34.58 | 31.25 |
| Latency to 5m | -0.51 | -0.76 |
| Latency to sing/duet | 0.58 | -0.54 |
| Time within 5m | 0.75 | 0.53 |
| Song/duet rate | -0.56 | 0.55 |
| Flyby rate | 0.51 | -0.31 |
| **(b) Male Principal Component Analysis** |  |  |
|  | PC1 | PC2 |
| Standard deviation | 1.98 | 1.46 |
| Proportion of Variance | 39.69 | 29.10 |
| Latency to 5m | -0.75 | -0.57 |
| Latency to sing/duet | 0.44 | -0.76 |
| Time within 5m | 0.83 | 0.24 |
| Song/duet rate | -0.32 | 0.70 |
| Flyby rate | 0.67 | -0.10 |

Pre-GnRH androgen levels were not explained by playback time (*p* = 0.89 in females and 0.19 in males), time-of-day bled (*p* = 0.93 in females and 0.93 in males), or net-to-bleed time (*p* = 0.62 in females and 0.62 in males), so we did not include these variables in final models. Likewise, post-GnRH androgens were not affected by playback time (*p* = 0.91 in females and 0.51 in males), time-of-day bled (*p* = 0.87 in females and 0.50 in males), or GnRH-to-bleed time (*p* = 0.31 in females and 0.52 in males) and were therefore excluded from analysis of STI response.

The final model for PC 1 STI response (rapid and long close approach and less vocal territoriality) in females included only a non-significant effect of pre-GnRH androgens (*p* = 0.15; Table S3a). Pre-GnRH androgens were a significant predictor (*p = 0.04*) of female PC 2 scores (quick and sustained vocal territoriality and close proximity), with higher androgen levels associated with reduced response to simulated intruders (Figure 2). No other predictors were included in the final model (Table S3b).

**Figure 2.**
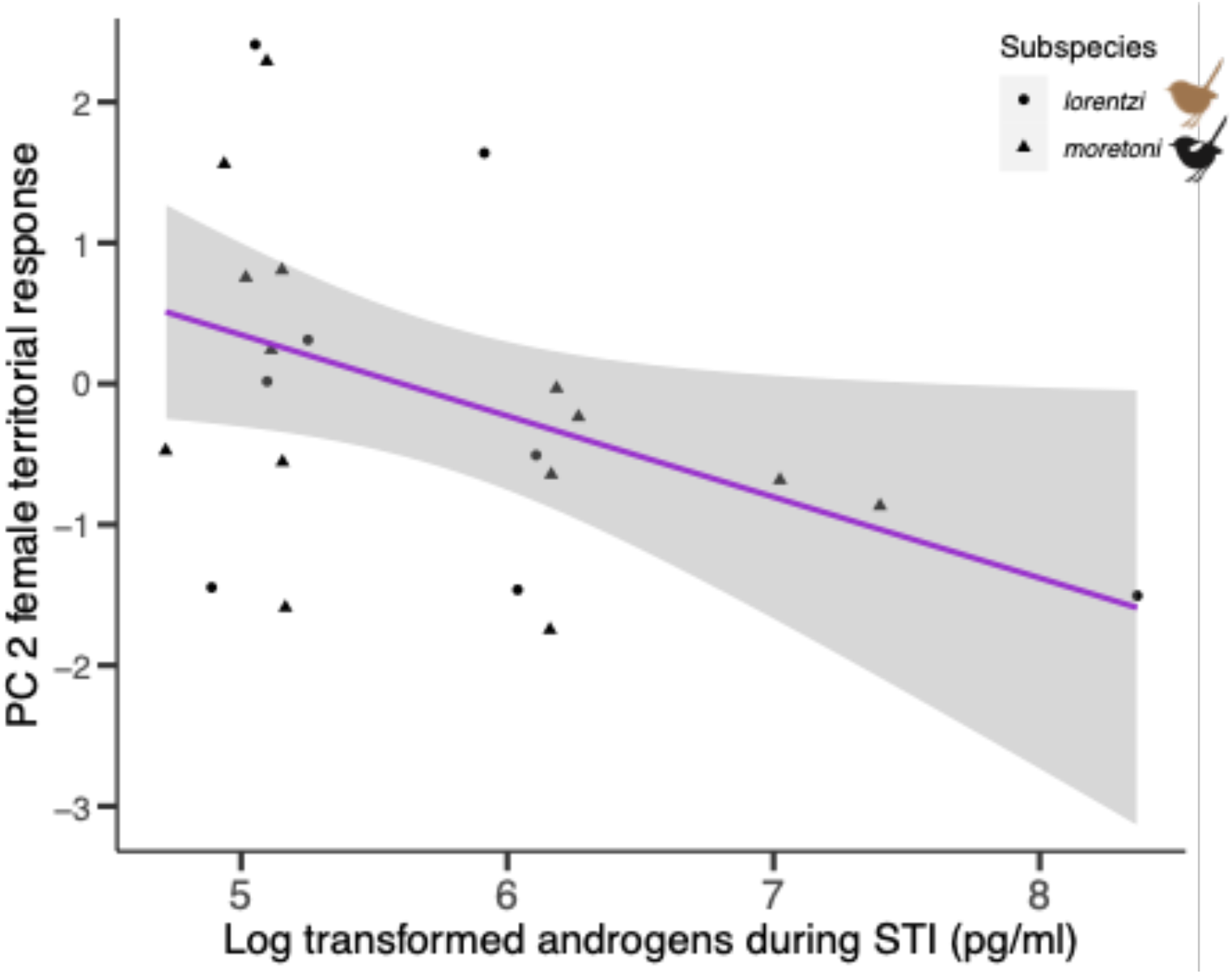
Log transformed androgens sampled during a simulated territorial intrusion (pg/ml) by principal component (PC) 2 from analysis of female territorial response. Log transformed (pre-GnRH) androgens sampled during STI were predictive of PC 2 scores in females (*p* = 0.04), which reflect shorter latency and greater rate of singing and shorter latency and longer duration of close approach to mounts. Subspecies differing in female ornamentation are separated by point shape.

For males, there were no significant predictors of PC 1 (rapid and sustained close approach and less vocal territoriality; Table S4a). There was a significant interaction between subspecies and pre-GnRH androgens on PC 2 (short latency and high rate of singing and rapid mount approach; *p* = 0.03; Table S4b), though neither subspecies showed a clear relationship between androgens and PC 2 scores (Figure S2). The interaction between subspecies and post-GnRH androgens was also included as a non-significant effect (*p* = 0.09).

### 3.3 Challenge hypothesis test in females

We compared androgen detectability and circulating androgen levels among 31 females captured during territorial defense to 20 flushed controls. The only significant predictor for whether androgens were detectable in our assay was breeding stage (Table S5a; *p* = 0.006). However, a post-hoc Tukey test did not reveal any significant differences among stages (laying/incubating vs. nestling/fledgling: *p* = 1.0, laying/incubating vs. not breeding: *p* = 0.12, nestling/fledgling vs. not breeding: *p* = 1.0). Log transformed pre-GnRH androgens did not differ among females sampled during STIs and flushed controls (Figure 3). Breeding stage (*p* = 0.10) and the interaction between sampling context and subspecies (*p* = 0.20) were non-significant predictors in the final model for pre-GnRH androgens (Table S5b). We used equivalence tests in both subspecies separately to test if STI and flushed androgen means were equivalent based on previously published differences in males from the same experiment (Boersma et al., 2022a). When subspecies were pooled, we found that mean androgens were equivalent across sampling contexts (*t* = 1.83, *p* = 0.04; mean ± sd: 522.42 ± 792.79 pg/ml during STI vs. 362.16 ± 259.05 pg/ml flushed). Given the interaction between subspecies and sampling context was included in the final model we also tested for equivalence in both subspecies separately. Mean androgens were not equivalent among STI and flushed sampling contexts in either *lorentzi* (*t* = -0.20, *p* = 0.58; mean ± sd: 647.96 ± 1139.63 pg/ml during STI vs. 213.15 ± 97.93 pg/ml flushed) or *moretoni* (*t* = -0.92, *p* = 0.18; 431.74 ± 414.27 pg/ml during STI vs. 461.50 ± 287.91 pg/ml flushed).

**Figure 3.**
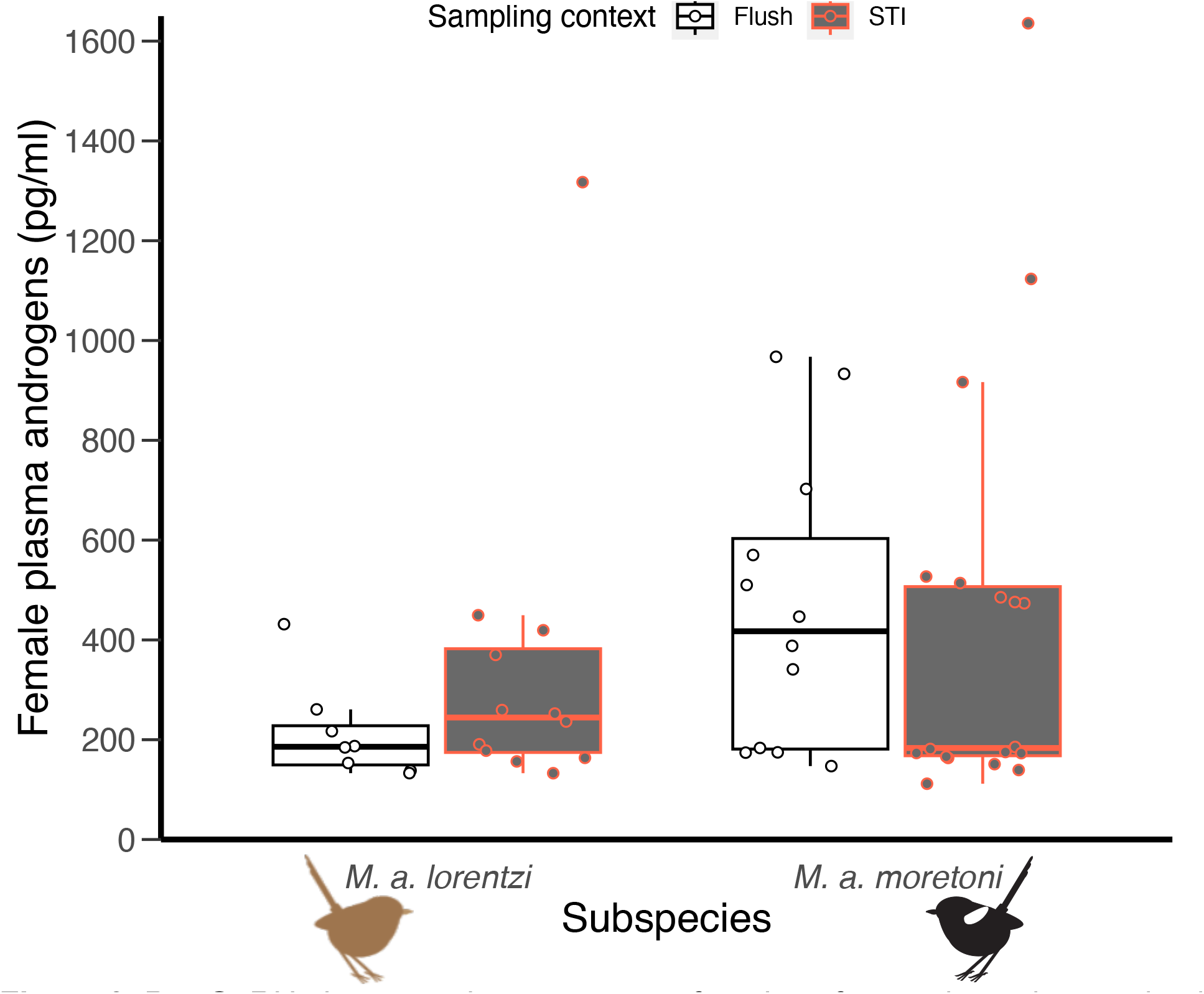
Pre-GnRH plasma androgens among females of two subspecies varying in ornamentation sampled by flushing or while responding to a simulated territorial intruder (STI). There were no differences between sampling contexts or subspecies. One *lorentzi* female sample during STI not shown due to falling out of plot area (4297.58 pg/ml).

## 4. Discussion

We tested whether differential capacity to produce androgens following GnRH challenge underlies previously found differences of mean androgen circulation, female ornamentation, and territoriality across subspecies of White-shouldered Fairywren (Table 1). Baseline androgens are higher on average in females of the mutually-ornamented and more territorial *moretoni* subspecies relative to unornamented *lorentzi* females (Enbody et al., 2018), whereas male androgens are highest in the more social *lorentzi*, despite a lack of variation in male ornamentation (Boersma et al., 2022a). A previous testosterone implant experiment showed that ornamentation can be induced in females of the *lorentzi* subspecies, which then caused an increase in territory defense behaviors (Boersma et al., 2020). In the current study, response to GnRH did not differ substantially across subspecies in either sex (Fig. 1), suggesting that subspecific variation in baseline androgens is not the product of an underlying capacity to produce and circulate androgens. Our null GnRH results across subspecies also indicate that capacity to elevate androgens does not underlie female phenotype variation. However, mean androgens following the GnRH challenge in females were not equivalent across subspecies, so it’s possible that small sample sizes constrained detecting significantly lower GnRH-induced androgens in unornamented *lorentzi*. Previous GnRH studies have found that capacity to produce androgens underlie variation in plumage phenotypes in male Dark-eyed Junco (McGlothlin et al., 2008) and White-throated Sparrow (Spinney et al., 2006).

Collectively, our null GnRH results and previous testosterone manipulation work might suggest that the lack of ornamentation observed in *lorentzi* females could be a result of constraints on androgen secretion— despite similar capacity to secrete androgens— that prevent elevation of androgens to the higher levels measured on average in *moretoni* (Enbody et al., 2018). If androgen secretion is constrained in *lorentzi* females, then we would expect that *lorentzi* females would show reduced androgenic response to social challenges like simulated territorial intrusions. However, neither subspecies elevated androgens significantly relative to flushed controls and mean androgens were equivalent across groups when subspecies were pooled. The highest androgens in this study were measured in female *lorentzi* sampled during STI (Figure 3), and mean differences between STI and flushed androgens were not statistically equivalent when analyzing *lorentzi* separately. It is possible that female plumage variation is the result of differences in response to circulating androgens across subspecies (e.g. receptors, enzymes, transcription factors, etc.(Bergeon Burns et al., 2014b; Béziers et al., 2017; Fuxjager et al., 2013; Hau and Goymann, 2015; Rosvall et al., 2012; Sewall, 2015; Soma et al., 2003). However, studies of gene expression in developing feather follicles of female White-shouldered Fairywrens suggest that these subspecies do not differ in the expression of androgen (or estrogen) receptors (Enbody et al., 2022), but receptor abundance may differ in other tissue.

Androgens like testosterone are known to mediate expression and degree of territorial behavior in males across diverse taxa (Hau et al., 2000; Oliveira et al., 2009; Pärn et al., 2008; Maria I Sandell, 2007; Sperry et al., 2010), and androgens can vary with territoriality in females (Cain and Ketterson, 2013, 2012; Kriner and Schwabl, 1991a; Pärn et al., 2008; Rosvall, 2013). In our study species, territory density and defense are higher on average in both sexes of *moretoni* relative to *lorentzi* (Enbody et al., 2019, 2018). We assessed whether individual and subspecific variation in response to simulated territorial intrusions were associated with higher pre-GnRH and GnRH-induced androgens, which would suggest a functional link between androgen secretion and/or capacity and territoriality. We did not find support for greater androgen secretion or capacity correlating with degree of territoriality. Pre-GnRH androgens sampled during an intrusion were somewhat predictive of response to simulated intruders in both sexes, though the direction of the effect was negative, which is surprising given that androgens are known to induce territory defense and aggressive behaviors across diverse taxa (Kriner and Schwabl, 1991b; Ros et al., 2002; Maria I. Sandell, 2007; Soma, 2006; Sperry et al., 2010). In females, pre-GnRH androgens were negatively associated with territorial response in the second principal component, corresponding to greater vocal territoriality and close approach to mounts (Fig. 2), which is consistent with previously observed territoriality in this system (Boersma et al., 2020; Enbody et al., 2018). The relationship between pre-GnRH androgens and territoriality was less clear in males, though we find a negative correlation between pre-GnRH androgens and PC 2 scores in *lorentzi* specifically. Therefore, we do not find evidence for elevated circulating androgens being a prerequisite for territory defense in this system. It’s possible that 10 minutes of playback time was insufficient for causing elevated androgens in this species, as another study found 30 minutes was required to elevate androgens (Wikelski et al., 1999). Though we did not find a relationship between playback time and androgen elevation in this experiment (Boersma et al., 2022a), only 6 individuals were sampled after 30 minutes.

No metric of response to GnRH challenge explained response to simulated territorial intrusions in either sex. Taken with the result that females did not substantially elevate androgens during territorial intrusions and a complementary study showing only marginally higher androgens in males during intrusions (Boersma et al., 2022a), we do not find that territory defense is contingent upon the capacity to elevate androgens. As with plumage, the higher territory defense in *M. moretoni* could be maintained and modulated by androgen receptor densities (Hau et al., 2004; Rosvall, 2013; Rosvall et al., 2012; Wingfield et al., 2001), or (unlike plumage) other sex steroids like progesterone (Adreani and Mentesana, 2018; Goymann et al., 2008), estradiol (Pärn et al., 2008; Soma et al., 2008, 2000), or dehydroepiandrosterone (DHEA, (Soma et al., 2015, 2002). These alternative mechanisms could allow maintenance of competitive behaviors absent costs of sustained androgen elevation (Wingfield et al., 2001).

Both subspecies of White-shouldered Fairywren tested in this study breed opportunistically, with both sexes showing breeding readiness year-round except for in cases of extreme drought (Boersma et al., 2022b); Enbody et al. 2019). Whereas most males responded to GnRH challenges by elevating androgens in relation to their pre-GnRH sample (N = 47 of 68 males, 69%), comparatively fewer females elevated androgens following GnRH challenge (N = 15 of 54 females, 28%). The low percentage of response to GnRH may suggest that many females reduce ovarian endocrine activity when they are not breeding. Though we did not detect a clear effect of breeding stage on whether a female elevated androgens following GnRH challenge, we used a coarse assessment of breeding stage based upon presence/absence of fledglings on territories and whether females were carrying eggs or had highly vascularized brood patches.

Therefore, a lack of effect of breeding stage on GnRH response should not be taken as conclusive evidence that capacity to elevate androgens does not differ across breeding stages, as has been shown in other species (George and Rosvall, 2018; Jawor et al., 2007). As a final consideration, we measured androgens in blood drawn from the jugular vein, which is thought to reflect steroid metabolism by the brain and brain-derived steroids (Newman et al., 2008; Saldanha and Schlinger, 1997; Soma et al., 2008). One might then suspect that we failed to measure androgens produced by the gonads in response to GnRH challenge. However, given that greater than two thirds of males increased androgens following injection, it seems unlikely that our jugular samples failed to capture gonad-derived androgens in response to GnRH. Whereas we did not include a vehicle-injected control group for our GnRH experiment, androgen concentrations typically decline following capture and handling (Lindsay, 2010; Vernasco et al., 2019; Woodley and Lacy, 2010) and the increases we measured in post-GnRH samples are likely to reflect natural variation in capacity to elevate androgens.

## Conclusions

In our study of White-shouldered Fairywrens, females subspecies that differ in baseline circulating androgens, territoriality, and ornamentation did not differ in androgen response to either GnRH or simulated intrusion challenges. Higher baseline androgens in males of the unornamented female *lorentzi* subspecies (Boersma et al., 2022a) coupled with our null GnRH results across male subspecies suggest that the ornamented female phenotype is not the product of correlated selection on androgen production in males. Collectively, studies of this species suggests that androgens have a role in activating female plumage ornamentation and enhanced territorial aggression underlying the ornamented phenotype (Boersma et al. 2020, Enbody et al. 2018).

Future work on androgen response pathways (i.e., receptors and enzymes) could resolve the full extent to which androgens mediate female ornamentation and behavior in this species. Determining the mechanisms underlying variation in female ornamentation and behavior will greatly improve our understanding how phenotypes evolve across diverse taxa.

## Acknowledgements

This work wouldn’t have been possible without the support of traditional landowners at each of our field sites in Papua New Guinea. Many individuals assisted with fieldwork, and we’d like to thank J. A. Gregg, D. Nason, M. Olesai, J. Sarusaruna, John, Kipling, Kingsford, and Manu for their expert help in the field.

## Supplementary materials

**Table S1.** Model terms for analysis of GnRH-induced plasma androgens in females. We analyzed three response variables: (a) a binary variable characterizing whether individuals responded to GnRH by elevating androgens, (b) the change (Δ) in androgens following GnRH administration, and (c) androgen titres post-GnRH among those that elevated androgens in response to challenge. All androgen measurements were log-transformed prior to analysis. Italicized rows depict predictors removed from final model, while bolded values show significant predictors.

(a) GnRH responders vs. non-responders (N = 54 females)
| Predictor | $\chi^2$ | DF | p |
| --- | --- | --- | --- |
| *Subspecies* | 0.59 | 1 | 0.44 |
| *Breeding stage* | 0.43 | 2 | 0.81 |
| **GnRH-to-bleed time** | **4.99** | **1** | **0.03** |
| *Time-of-day bled* | 0.04 | 1 | 0.83 |

(b) $\Delta$ Androgens post-GnRH (N = 53 females)
| Predictor | SS | DF | F | p |
| --- | --- | --- | --- | --- |
| *Subspecies* | 0.28 | 1 | 0.50 | 0.48 |
| *Breeding stage* | 1.53 | 2 | 1.37 | 0.26 |
| *GnRH-to-bleed time* | 0.17 | 1 | 0.29 | 0.59 |
| *Time-of-day bled* | 1.79 | 1 | 3.11 | 0.08 |

(c) GnRH-induced androgens (N = 14 females)
| Predictor | SS | DF | F | p |
| --- | --- | --- | --- | --- |
| *Subspecies* | 0.43 | 1 | 1.14 | 0.31 |
| *Breeding stage* | 0.15 | 2 | 0.14 | 0.87 |
| *GnRH-to-bleed time* | 0.16 | 1 | 0.39 | 0.55 |
| **Time-of-day bled** | **4.26** | **1** | **12.16** | **0.004** |

**Table S2.**
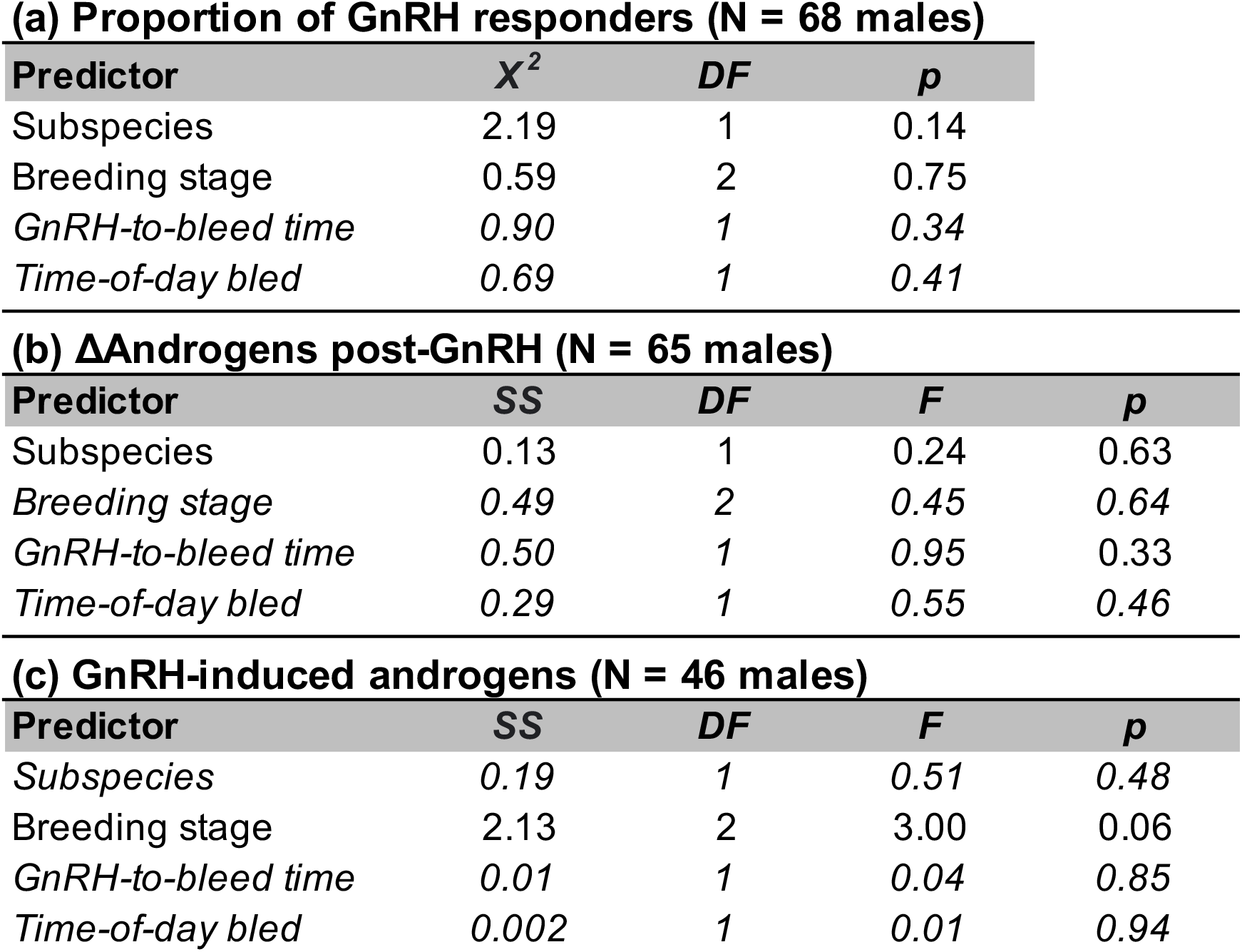
Model terms for analysis of GnRH-induced plasma androgens in males. We analyzed three response variables: (a) a binary variable characterizing whether individuals responded to GnRH by elevating androgens, (b) Δ androgens following GnRH administration, and (c) androgen titres post-GnRH among those that elevated androgens in response to GnRH challenge. All androgen measurements were log-transformed prior to analysis. Italicized rows depict predictors removed from final model, while bolded values show significant predictors.

**Table S3.** Model parameters for predictors of principal components from female simulated territorial intrusion analysis. Italicized rows depict predictors removed from final model, while bolded values show significant predictors.

**(a) Principal component 1 (N = 22 females)**
| **Predictor** | **SS** | **DF** | **F** | **p** |
| --- | --- | --- | --- | --- |
| pre-GnRH androgens | 4.25 | 1 | 2.21 | 0.15 |
| *post-GnRH androgens* | *0.02* | *1* | *0.01* | *0.92* |
| *Subspecies* | *0.63* | *1* | *0.30* | *0.59* |
| *Subspecies\*pre-GnRH androgens* | *0.06* | *1* | *0.03* | *0.86* |
| *Subspecies\*post-GnRH androgens* | *2.07* | *1* | *0.98* | *0.34* |
| *Breeding stage* | *6.13* | *2* | *1.62* | *0.23* |

**(b) Principal component 2 (N = 22 females)**
| **Predictor** | **SS** | **DF** | **F** | **p** |
| --- | --- | --- | --- | --- |
| ***pre-GnRH androgens*** | **6.06** | **1** | **4.68** | **0.04** |
| *post-GnRH androgens* | *0.27* | *1* | *0.20* | *0.66* |
| *Subspecies* | *0.03* | *1* | *0.02* | *0.89* |
| *Subspecies\*pre-GnRH androgens* | *0.003* | *1* | *0.002* | *0.97* |
| *Subspecies\*post-GnRH androgens* | *0.14* | *1* | *0.09* | *0.76* |
| *Breeding stage* | *2.51* | *2* | *0.98* | *0.40* |

**Table S4.** Model parameters for predictors of principal components from male simulated territorial intrusion analysis. Italicized rows depict predictors removed from final model, while bolded values show significant predictors.

**(a) Principal component 1 (N = 17 males)**
| **Predictor** | **SS** | **DF** | **F** | **p** |
| --- | --- | --- | --- | --- |
| *pre-GnRH androgens* | *1.48* | *1* | *0.86* | *0.37* |
| *post-GnRH androgens* | *0.0003* | *1* | *0.0002* | *0.99* |
| *Subspecies* | *0.02* | *1* | *0.01* | *0.92* |
| *Subspecies\*pre-GnRH androgens* | *0.02* | *1* | *0.01* | *0.92* |
| *Subspecies\*post-GnRH androgens* | *1.29* | *1* | *0.62* | *0.45* |
| *Breeding stage* | *3.97* | *2* | *1.16* | *0.34* |

**(b) Principal component 2 (N = 17 males)**
| **Predictor** | **SS** | **DF** | **F** | **p** |
| --- | --- | --- | --- | --- |
| *pre-GnRH androgens* | *0.63* | *1* | *0.53* | *0.48* |
| *post-GnRH androgens* | *0.23* | *1* | *0.19* | *0.67* |
| *Subspecies* | *0.07* | *1* | *0.06* | *0.81* |
| ***Subspecies\*pre-GnRH androgens*** | **7.46** | **1** | **6.24** | **0.03** |
| *Subspecies\*post-GnRH androgens* | *4.25* | *1* | *3.56* | *0.09* |
| *Breeding stage* | *1.96* | *2* | *0.74* | *0.50* |

**Table S5.** Model parameters for challenge hypothesis test in females. Sampling context refers to simulated territorial intrusion (STI) vs. flushed controls. Italicized rows depict predictors removed from final model.

**(a) Androgen detectability (N = 51 females)**
| Predictor | $\chi^2$ | DF | p |
| --- | --- | --- | --- |
| *Sampling context* | 0.47 | 1 | 0.49 |
| *Subspecies* | 0.11 | 1 | 0.74 |
| *Sampling context\*Subspecies* | 0.69 | 1 | 0.41 |
| **Breeding stage** | **10.24** | **2** | **0.006** |
| *Net-to-bleed time* | 0.48 | 1 | 0.49 |
| *Time-of-day bled* | 0.65 | 1 | 0.42 |

**(b) pre-GnRH plasma androgens (N = 51 females)**
| Predictor | SS | DF | F | p |
| --- | --- | --- | --- | --- |
| Sampling context | 0.09 | 1 | 0.15 | 0.70 |
| Subspecies | 0.47 | 1 | 0.80 | 0.38 |
| Sampling context*Subspecies | 1.01 | 1 | 1.71 | 0.20 |
| Breeding stage | 2.91 | 2 | 2.47 | 0.10 |
| *Net-to-bleed time* | 0.25 | 1 | 0.42 | 0.52 |
| *Time-of-day bled* | 0.13 | 1 | 0.21 | 0.65 |

**Figure S1.**
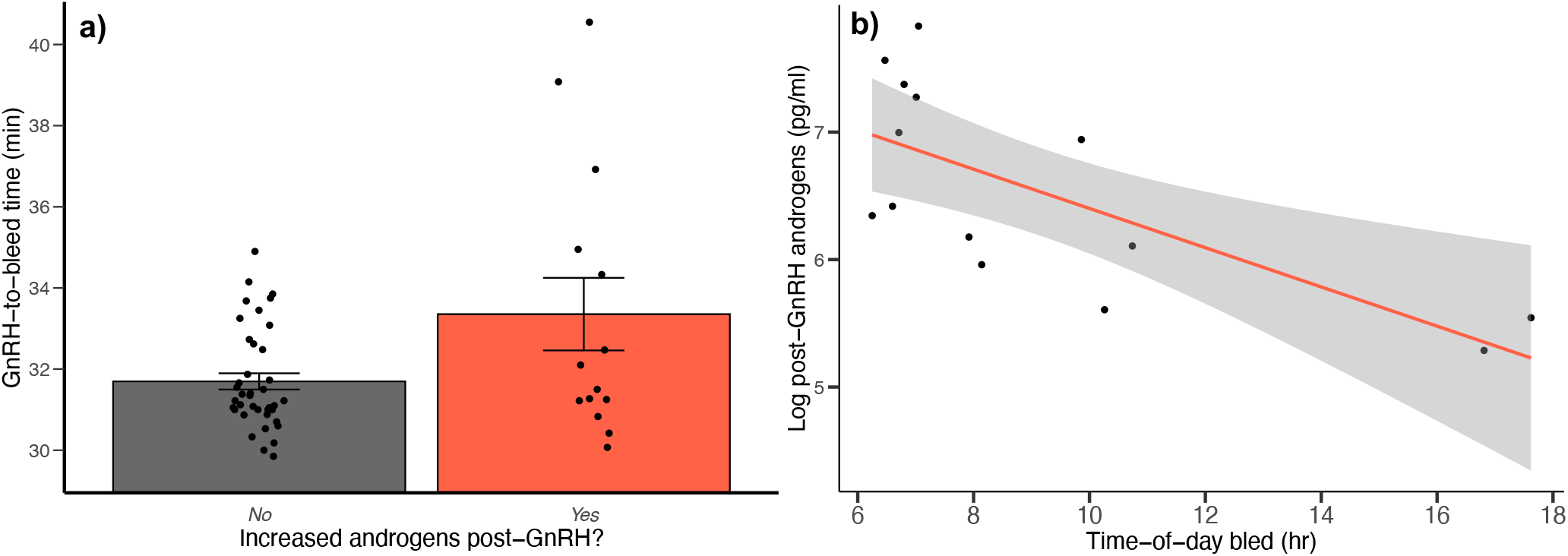
Relationship between GnRH-to-bleed time and whether females increased androgens following GnRH challenge **(a)** and effect of time-of-day bled on log transformed post-GnRH androgens in females **(b)**.

**Figure S2.**
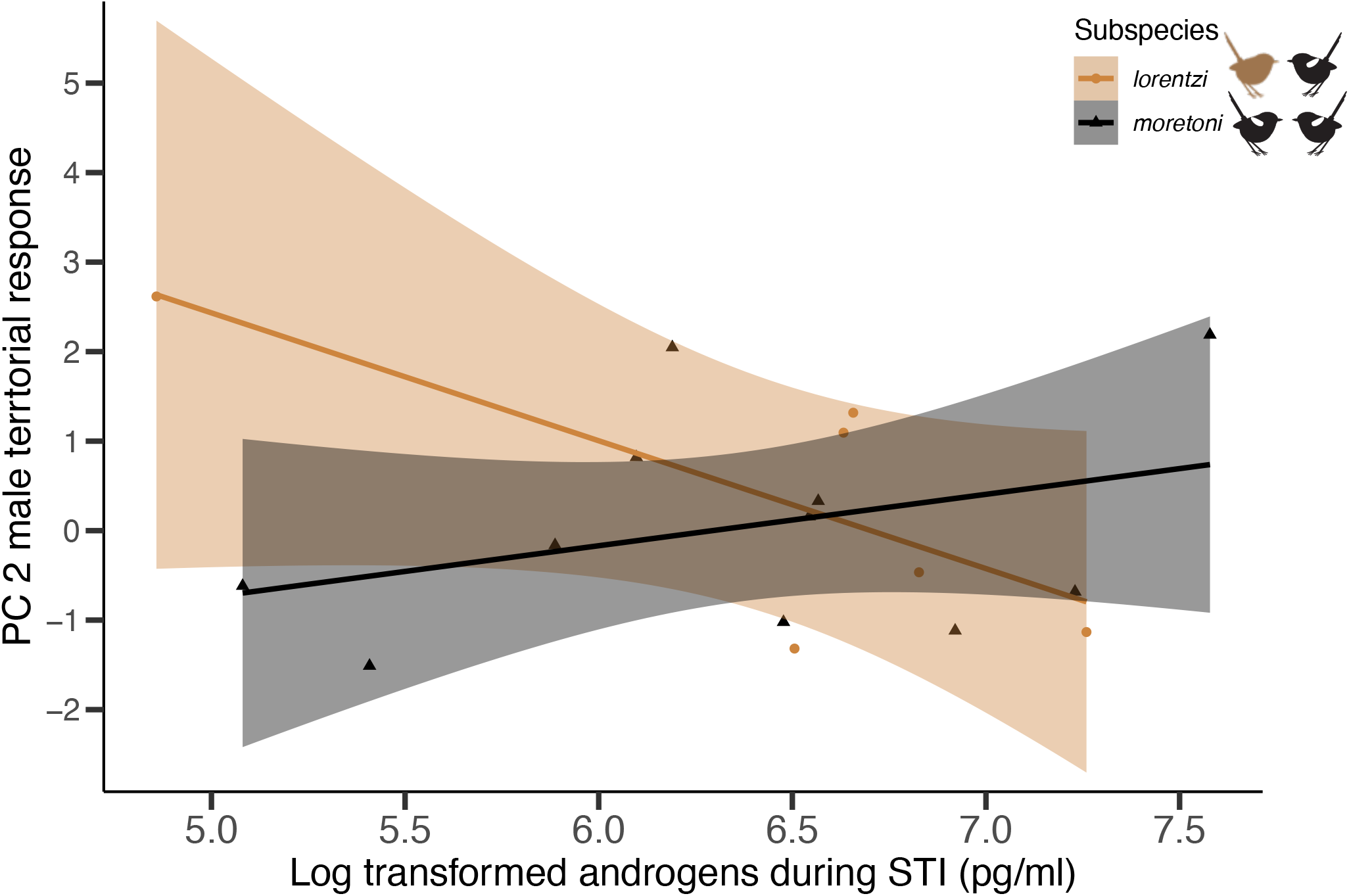
Pre-GnRH plasma androgen residuals (accounting for capture parameters) and principal component (PC) 2 of male response to simulated territorial intrusion among two subspecies. There was a significant interaction between subspecies and pre-GnRH androgens on PC 2 (*p* = 0.04).

